# Ancestral role of Fat-like cadherins in planar cell polarity

**DOI:** 10.1101/602029

**Authors:** Maria Brooun, Alexander Klimovich, Mikhail Bashkurov, Bret J. Pearson, Robert E. Steele, Helen McNeill

**Affiliations:** Lunenfeld-Tanenbaum Research Institute, Toronto, ON, Canada; Zoological Institute, Christian-Albrechts University of Kiel, Germany; The Hospital for Sick Children, Program in Developmental and Stem Cell Biology, Toronto, ON, Canada; University of Toronto, Department of Molecular Genetics, Toronto, ON, Canada; Department of Biological Chemistry, University of California, Irvine, CA, USA; Department of Developmental Biology, Washington University School of Medicine, St. Louis, MO, USA

**Author notes:** Correspondence should be addressed to: Helen McNeill or Maria Brooun. Lead Author: Helen McNeill.

**Keywords:** *Hydra*, cadherins, planar cell polarity, regeneration

## Abstract

Fat family cadherins are enormous proteins that regulate planar cell polarity (PCP) and cell adhesion in bilaterian animals. Their evolutionary origin can be traced back to prebilaterian species, but their ancestral function(s) are unknown. We identified Fat-like and Dachsous cadherins in *Hydra*, a member of the early-diverging metazoan phylum Cnidaria. *Hydra* has a simple body plan with only two epithelial layers and radial symmetry. We find that *Hydra* homologues of Fat-like (HyFat) and Dachsous (HyDs) co-localize at the apico-lateral membrane of ectodermal epithelial cells. Remarkably, HyFat is planar polarized perpendicular to the oral-aboral axis of the animal. Using knockdown approaches we found that HyFat is involved in the regulation of local cell alignment, but is dispensable for the global alignment of ectodermal myonemes along the oral-aboral axis. The intracellular domain (ICD) of HyFat is involved in the morphogenesis of ectodermal myonemes. Thus, Fat family cadherins have ancient, prebilaterian functions in cell adhesion, tissue organization and planar polarity.

## INTRODUCTION

Planar cell polarity (PCP) describes the coordinate polarization of cells within a tissue. PCP regulates a wide range of organization in animal development, from hairs on insect wings, to convergent extension movements during vertebrate gastrulation, and oriented cell divisions across species [1]. The mechanisms controlling PCP were first deciphered in *Drosophila*, and have been shown to be conserved in bilaterians. Two modules are known to regulate PCP. The most studied and best understood is known as the core module, consisting of a number of transmembrane proteins and associated cytoplasmic components, such as Frizzled/Disheveled and Strabismus/Prickle (reviewed in [1]).

Another critical regulator of PCP is the Fat/Dachsous (Fat/Ds) complex, consisting of two giant atypical cadherins, Fat and Dachsous. Fat and Dachsous interact via extracellular cadherin domains, and use their intracellular domains to transduce planar polarity signals within the cell, as well as to control proliferation via the Hippo pathway and mediate metabolic control (reviewed in [2]). In addition, a closely related family of Fat-like cadherins, similar in their extracellular domains but with a distinct intracellular domain, can polarize cells by influencing actin and microtubule dynamics [3–6].

Fundamental questions regarding the origin and ancestral functions of PCP genes remain unanswered. How did the PCP machinery evolve, and what is the minimal machinery required to polarize a cell within a plane? Homologues of PCP proteins have been identified across the animal kingdom [7, 8], however, functional studies in non-bilaterian metazoans are very limited. Only the core PCP proteins have been studied in non-bilaterians. For example, Strabismus is required for gastrulation in the cnidarians *Nematostella* and *Clytia* [9, 10]. Core PCP proteins have also been identified in the cnidarian *Hydra* [11], however, the presence and function of Fat or Fat-like cadherins in cnidarians have not been explored. The freshwater cnidarian polyp *Hydra* is radially symmetric and has only two cell layers, an ectoderm and an endoderm. Both layers are composed of epithelial cells with intermingled cells of the interstitial stem cell lineage [12]. The *Hydra* body column is a cylinder, with a head at the apical (oral) end and a foot at the basal (aboral) end (Figures 1A,B). The head contains the mouth and a ring of tentacles. Epithelial cells of the body column divide continuously and are constantly displaced towards the oral and aboral ends where they differentiate into head and foot-specific cells and are eventually sloughed off [13]. Epithelial cells of both layers extend muscle-like processes called myonemes that are oriented along the oral/aboral axis in the ectoderm and circumferentially in the endoderm. The direction of cell displacement along the oral-aboral axis changes when cells enter the tentacles and when a new bud, *Hydra’s* asexual form of reproduction, is formed (Figures 1A,B) [14].

**Figure 1.**
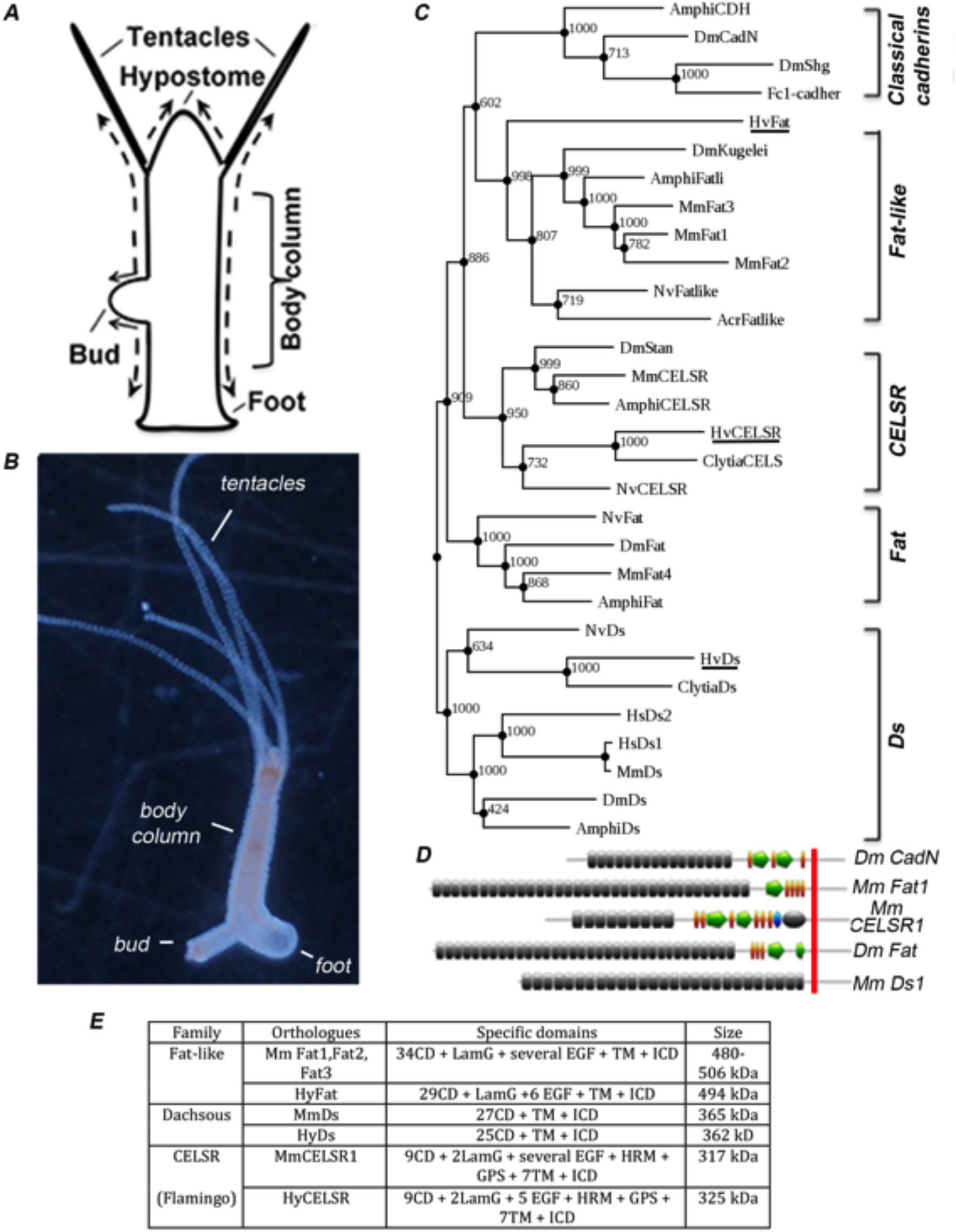
Cnidarian *Hydra* possess single homologues of atypical cadherins Fat-like, Dachsous and CELSR. (A) Schematic of the *Hydra* body plan, arrows indicate directions of cell displacement; (B) Budding adult *Hydra* polyp; (C) Phylogenetic tree of Fat, Fat-like, Dachsous and CELSR cadherin subfamilies based on the sequences of all cadherin domains, Maximum Likelihood analysis (1000 bootstrap replicates, bootstrap values are indicated for each node) reveals the clustering of predicted *Hydra* cadherin homologues with the corresponding protein subfamilies; *Hydra vulgaris* homologues are underlined; (D) Domain composition of representative atypical cadherins and classical cadherin, 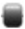 - cadherin domain, 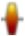 - EGF repeat-like domain, 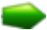 - laminin domain, 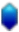 - hormone receptor domain, 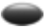 - latrophilin/CL-1-like GPS domain, 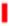 - cellular membrane; (E) Table showing the domain composition of atypical mammalian cadherins and their *Hydra* homologues, domain abbreviations: CD – cadherin domain, LamG – laminin G domain, EFG – epidermal growth factor-like domain, TM – transmembrane domain, ICD – intracellular domain, HRM – hormone receptor domain, GPS – latrophilin/CL-1-like GPCR proteolytic site domain.

In *Hydra* expression of the core PCP genes *frizzled* and *dishevelled* is strongly upregulated at the bases of the tentacles and early in bud evagination [11], suggesting the core PCP module is acting in those locations. However, expression of core PCP genes along the body column, where cells exhibit planar polarization along the oral-aboral axis is weak, suggesting another PCP mechanism could be acting there.

Here we show for the first time that a Fat-like cadherin regulates tissue organization in a non-bilaterian animal. We found that HyFat is apically localized in epithelial cells and planar polarized along the oral-aboral axis, indicating that the polarized location of Fat-like cadherins evolved prior to the divergence of cnidarians and bilaterians. Overexpression experiments suggest that the intracellular domain of HyFat is involved in the morphogenesis of the myonemes. Transgenic knockdown revealed that HyFat has a critical role in tissue level organization of the body column.

## RESULTS

### *Hydra* Fat-like and Dachsous are expressed in epithelial cells

Our analysis of *Hydra* genomic, transcriptomic, and EST data identified a single *Fat-like* cadherin gene (*HyFat*) and a single *Dachsous* gene (*HyDs*). HyFat and HyDs share a similar protein structure with their vertebrate homologues (Figure 1E). However, the intracellular domains (ICDs) of HyFat and HyDs show only low similarity with *Drosophila* and vertebrate homologues, and much higher similarity with homologues from the medusozoan group of cnidarians (Figures S1A,B). HyFat ICD shares an HWD motif with mammalian Fat1 and Fat3 (Figure S1A). This domain is present in ICDs of most Fat-like proteins, but not in ICDs of Fat, Dachsous, CELSR, or classical cadherin protein families (data not shown). In addition the ICDs of the Fat-like proteins of cnidarians have three proline-rich motifs (Figure S1B), which have similarity to EVH1-binding domains of MmFat1 [6, 15], suggesting a possible role for HyFat in the regulation of actin polymerization.

The conservation of the ICDs of bilaterian Ds family proteins is clearest among three homology regions [16], however, these regions are not present in ICD of HyDs nor Ds homologs of the medusozoans *Clytia, Alatina*, or *Cassiopea*, (Figure S1C). ICDs of those medusozoans, except that of *Hydra*, have two regions of homology with *Nematostella vectensis* Ds (NvDs). The ICD of NvDs, in turn, shares characteristic conserved regions with the other members of Ds family (Figure S1C). Phylogenetic analyses using either sequences comprising all cadherin domains or ICDs shows that the *Hydra* cadherins we identified belong to Fat-like and Dachsous families (Figures 1C, S2). A true Fat homologue is present in the anthozoan cnidarians *Nematostella* and *Acropora* [17]. However, we were unable to identify a homologue of Fat in the genomes of *Hydra* or other medusozoan cnidarians. Our *in silico* analysis also reveals a single *Hydra* homolog of Flamingo/ CELSR, another atypical cadherin involved in PCP [18], that we call HyCelsr (Figure S1D).

*In situ* hybridization shows that in adult *Hydra* polyps *HyFat* and *HyDs* are expressed in both ecto- and endodermal epithelial cells throughout the body column, with weaker expression in the tentacles, foot, and hypostome (Figures 2A,B, S3A,B). *HyFat* expression is elevated in the lower portion of the body column, just above the foot, and around the tentacle bases (Figure 2A). In conclusion, in Hydra there is a single Fat-like (HyFat) and Ds (HyDs) that are expressed in epithelial cell lineages.

**Figure 2.**
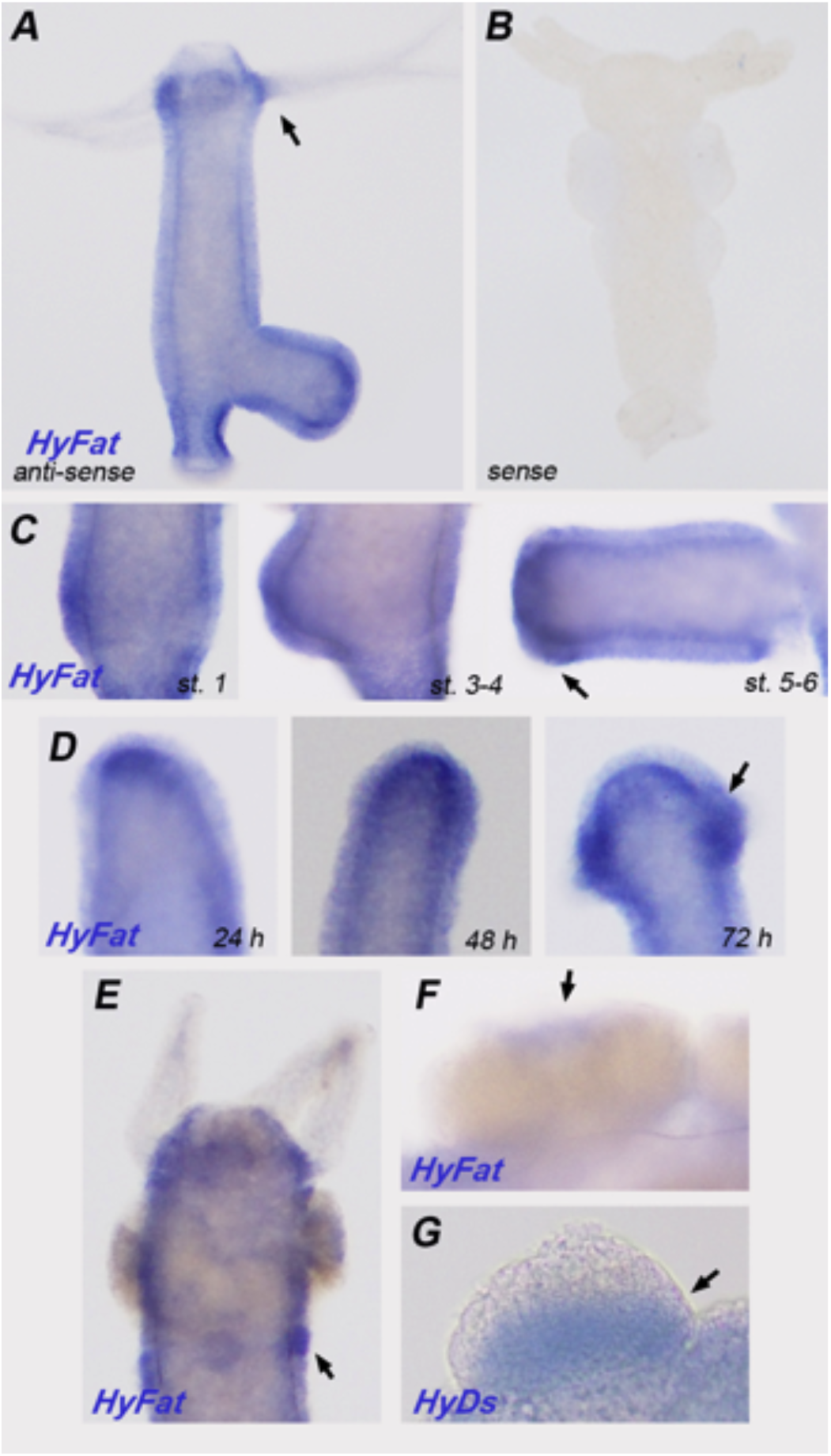
*HyFat* is expressed throughout the body column and upregulated during head and testis development. (A,B) HyFat is expressed throughout the animal; (C) Upregulation of *HyFat* during budding (budding stages as in [21]); (D) Expression of *HyFat* in a regenerating head, times after bisection are indicated; arrows indicate the expression of *HyFat* around the tentacles and at the presumptive tentacle zone. (E,F,G) Expression of HyFat and HyDs in early (E) and mature (F,G) testes, arrows indicate an upregulation of gene expression within the testes.

### Upregulation of *HyFat* during head formation and spermatogenesis

The *Hydra* head can form *de novo* during bud development [14], and during regeneration. Budding starts as a thickening of the ectoderm by an evagination of both layers [21]. The bud develops into a small polyp with a head at the distal end. The bud detaches after formation of a basal disk at the proximal end. Upregulation of *HyFat* was observed at the tip of a young bud and in the presumptive tentacle area in very early buds (Figure 2C). The elevated expression of *HyFat* around the tentacles was maintained in the adult. During head regeneration upregulation of *HyFat* expression was seen first in the endoderm at the tip of a regenerating head, and then in the ectoderm of the tentacle buds (Figure 2D). This raises the possibility that *HyFat* plays a role in head formation.

During spermatogenesis in *Hydra*, spermatogonia, which are derived from a sperm-committed stem cell in the interstitial cell lineage, accumulate between the mesoglea and the ectoderm causing the ectodermal epithelium to form a testis [22, 23]. *HyFat* gene expression is upregulated in the ectoderm of the testis, first at the very early stage and then at the distal tip of the mature testis (Figures 2E,F). *HyDs* is also expressed during testis development. However, unlike *HyFat*, the upregulation of *HyDs* gene expression is observed not in the epithelial cells but within the spermatogonia (Figure 2G). *HyDs* gene expression is stronger in the proximal part of the testis and fades towards the tip. This complementary pattern of expression of *HyFat* and *HyDs* is notable, as mammalian Fat and Dachsous proteins are also expressed in a reciprocal pattern in the mouse embryonic hindbrain [24] and suggests a possible interaction of these two proteins along the boundaries of their expression. Interestingly, ectodermal epithelial cells divide the testis into compartments with long cytoplasmic strands, and this compartmentalization is believed to be essential for proper spermatogenesis [25].

*HyFat* gene expression is upregulated in the ectoderm of the testis, first at the very early stage and then at the distal tip of the mature testis (Figures 2E,F). *HyDs* is also expressed during testis development. However, unlike *HyFat*, the upregulation of *HyDs* gene expression is observed not in the epithelial cells but within the spermatogonia (Figure 2G). *HyDs* gene expression is stronger in the proximal part of the testis and fades towards the tip. This complementary pattern of expression of *HyFat* and *HyDs* is notable, as mammalian Fat and Dachsous proteins are also expressed in a reciprocal pattern in the mouse embryonic hindbrain [24] and suggests a possible interaction of these two proteins along the boundaries of their expression. Ectodermal epithelial cells divide the testis into compartments with long cytoplasmic strands, and this compartmentalization is believed to be essential for proper spermatogenesis [25].

### HyFat is planar-polarized at apical junctions in the body column

We generated polyclonal antibodies against the HyFat ICD (residues 4191 to 4393). Immunoblotting of *Hydra* lysates with anti-HyFat antiserum identified a polypeptide of 490 kDa, corresponding to the predicated molecular weight of full length HyFat. In addition, polypeptides of 180 kDa, 100 kDa and 35 kDa were specifically recognized by this antiserum (Figures 3F, 4C). Fat and Fat-like cadherins have been shown to be post-translationally cleaved [26, 27]; HyFat may also undergo such proteolytic processing.

**Figure 3.**
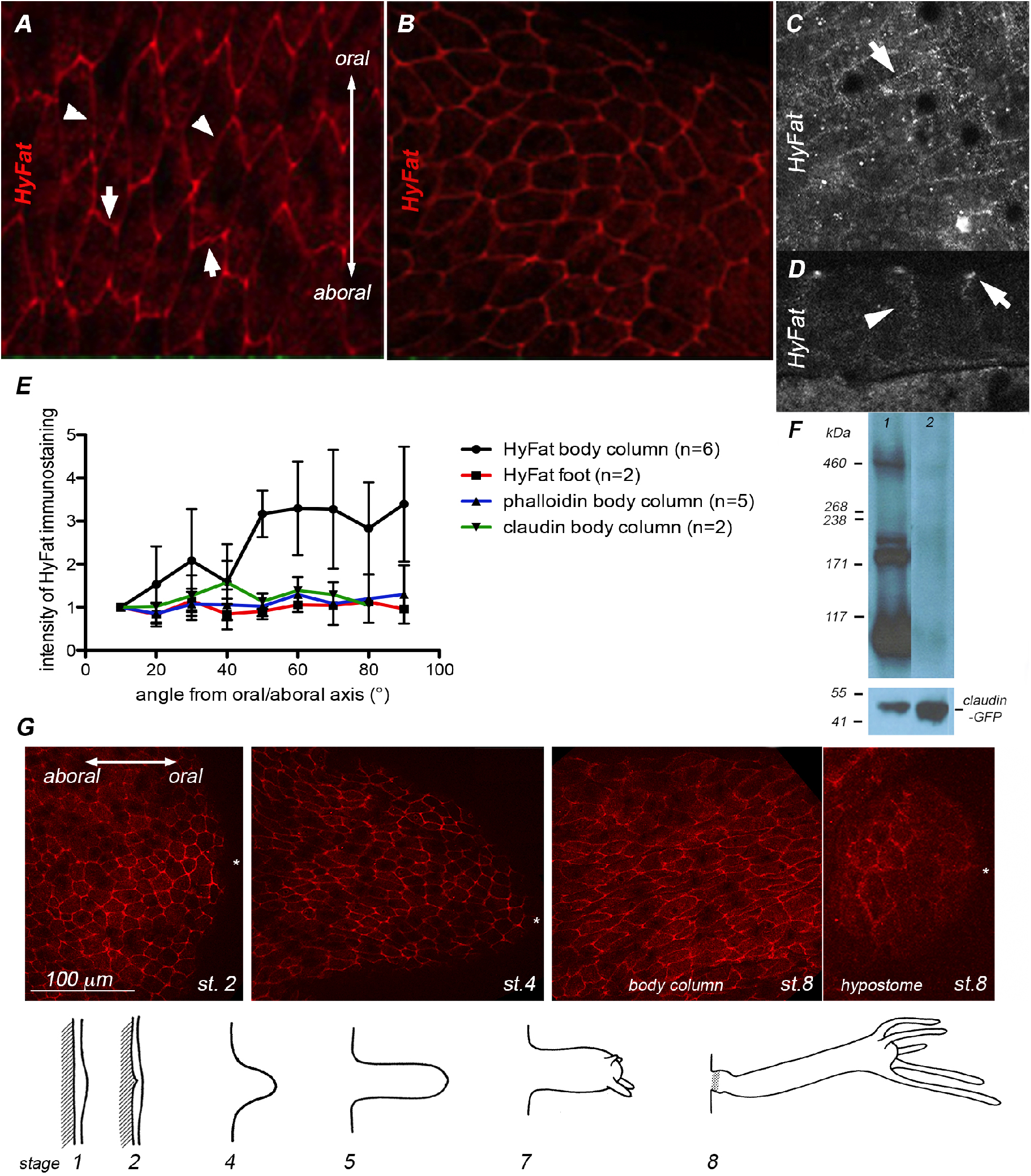
HyFat is expressed in *Hydra* epithelium and planar polarized in the ectoderm. Immunostaining with HyFat antiserum: (A) In the body column, HyFat is located at the junctions between ectodermal epithelial cells and, specifically, at the tricellular junctions; arrows indicate an enrichment of HyFat at the cell membrane perpendicular to the oral-aboral axis, arrowheads point to the cell membranes that are parallel to the oral-aboral axis; (B) The head; (C) Immunostaining of the endoderm with affinity-purified HyFat antibodies (1:100), arrowheads indicate the localization of HyFat at the cellular membrane; (D) Accumulation of HyFat at the apical (arrow) and lateral (arrowhead) cell membranes of the ectodermal epithelial cells; (E) Graph shows the polarized accumulation of HyFat along the apical membranes of ectodermal epithelial cells (n indicates number of experiments, 60-70 measurements for each experiment), the intensity of HyFat staining was normalized for the intensity of the membranes that are parallel (0-10°) to the oral/aboral axis. Data represent the mean with SD; (F) Immunoblot analysis of a total protein lysate from a transgenic *Hydra* line constitutively expressing claudin-GFP, HyFat anti-serum (1:1000) was preabsorbed with either GST (lane 1) or GST-HyFat antigen (lane 2); the lanes shown were selected from the original x-ray image; (G) Dynamic of HyFat localization during bud development, asterisk indicates the tip of the bud, budding schematic is adapted from [21].

Immunostaining of fixed *Hydra* polyps with HyFat antiserum and affinity purified anti-HyFat antibodies showed localization of HyFat at the apical and lateral ectodermal epithelial junctions (Figures 3A,B,D). This staining was especially strong at the tri-cellular apical junctions. Weaker HyFat staining was also observed at the endodermal epithelial membranes (Figure 3C). In the ectoderm, immunostaining of the apico-lateral membranes that are perpendicular to the axis of the animal appeared stronger than the staining of the membranes that are parallel to the axis (Figure 3A). Interestingly, this polarized localization of HyFat was observed in the body column, but not in the head or foot regions where the apical surfaces of ectodermal epithelial cells have a more rounded morphology (Figures 3A,B,E).

During bud development epithelial cells of the parent change their orientation and align along a new oral-aboral axis [28]. Early in bud development, ectodermal epithelial cells of the body column that are recruited into the bud lose planar polarization of HyFat (Figure 3G). As a bud develops, HyFat remains unpolarized in the future head cells, whereas in the cells of the future body column and cells that are being displaced from the parent into the bud, HyFat is polarized along the bud’s oral-aboral axis. Thus, HyFat appears to be planar polarized in the ectodermal epithelial cells that undergo both oriented cell division and oriented displacement.

We also generated antiserum against the ICD of HyDs (residues 2958 – 3113). Immunostaining with an affinity-purified HyDs antibody overlapped with HyFat staining at the apical junctions of ectodermal epithelial cells (Figure S3C,D), although no planar polarization was detected. Specificity of both anti-HyFat and anti-HyDs antibodies was demonstrated by preabsorption with a corresponding antigen (Figures S3C,D; S4). Thus, *Hydra* homologues of Fat-like and Dachsous cadherins localize to the apical junctions of ectodermal epithelial cells.

### Downregulation of *HyFat* interferes with local alignment and size of epithelial cells

To investigate the function of *HyFat*, we used an RNA hairpin approach [29]. Two different hairpin-expressing plasmid constructs, *shFat9* (nucleotides 12573 to 13178) and *shFat4* (nucleotides 11493 to 11792) were inserted downstream of GFP under the control of an actin gene promoter that is active in all epithelial cells [30]. Transgenic *Hydra* were obtained by injecting plasmid DNAs into 1-4 cell stage embryos [31]. We could not obtain stable transgenic lines expressing the hairpins under control of the actin promoter because hatchlings rapidly lost the transgenic cells. These data suggest that the downregulation of *HyFat* caused by constitutive expression of the hairpins led to loss of cells. One line bearing the *GFP-shFat4* survived for a month. To overcome the lethality of *shFat* knockdown, we tried doxycycline-dependent transcriptional activation. We cloned *shFat9* hairpin into the doxycycline-inducible vector pIndGFP, described in a previous study (Figure 4F) [30]. We were able to establish one *Hydra* line expressing *shFat9* hairpin. However, we found that *shFat9* hairpin was expressed constitutively, without induction by doxycycline (Figure S5), in both ecto- and endoderm (RT-PCR data not shown). *In situ* hybridization, immunoblotting, and immunostaining showed a dramatic decrease in the amount of *HyFat* transcript and HyFat protein in *shFat9* polyps (Figures 4C-H). Apical HyFat staining is weaker in *shFat9*, and staining of the lateral membranes is absent (Figures 4G,H). Immunostaining of *shFat9* with HyFat antiserum showed reduced levels of HyFat along the membranes of ectodermal epithelial cells and at the tricellular junctions (Figures. 4G-K). That the knockdown of *HyFat* in *shFat9* polyps is incomplete is evident from both immunoblotting and immunostaining.

**Figure 4.**
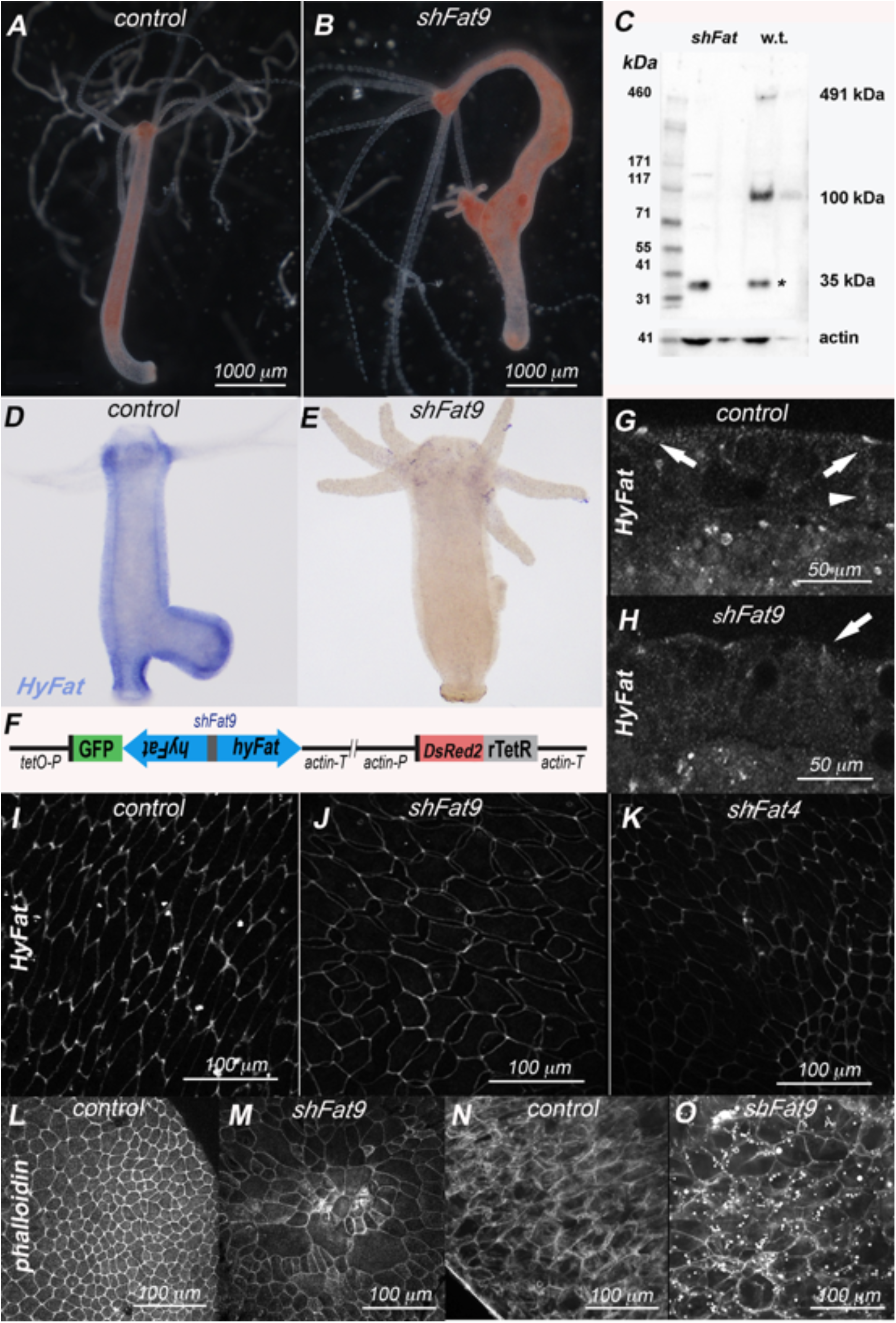
Downregulation of *HyFat* affects epithelial integrity and cell morphology. (A, B) *shFat9* mutant has an altered body column shape; (C) Immunoblot analysis of lysate from wild type and *shFat9*, asterisk indicates a non-specific 35 kDa band. (D,E) *In situ* hybridization *HyFat* antisense probe; (F) Schematic drawing of inducible *shFat9* construc (G,H) Decreased HyFat at the apical (arrow) and lateral (arrowhead) epithelial junctions in *shFat9* polyp; (I,J,K)Immunostaining with anti-HyFat antibodies reveals weaker adhesive contacts and irregularly shaped ectodermal cells in *shFat9* polyps; (L,M) Apical view of the ectodermal surface of the basal disk stained with phalloidin; (N,O) Confocal imaging of the endoderm stained with phalloidin.

Analysis of the *shFat9* partial knockdown animals revealed several phenotypic features. The body column of *shFat9* was notably elongated compared with wild type and had a ‘lumpy sock’ shape (Figures 4A,B). The body column of *shFat9 Hydra* resembled the body column of *Hydra* treated with1,2-diacyl-glycerol, which causes an elongation of the body column and formation of ectopic tentacles and heads [32]. However, ectopic tentacles were rare in *shFat9*, and we did not observe the formation of ectopic heads in *shFat9* animals. We did not observe an altered body shape with *shFat4*, probably due to a smaller number of transgenic cells. The apical contacts between ectodermal epithelial cells of *shFat9* and *shFat4* often become separated upon fixation (Figure 4J), which suggests an adhesion function for HyFat. Although the *shFat9* line was more stable than *shFat4*, elimination of transgenic cells occurred in *shFat9* polyps as well: we often observed multicellular junctions resembling healing wounds along the body column of *shFat9* animals (Figure S6). We propose that they are the result of healing at sites where cells have been lost. Over time, the *shFat9* line was overtaken by cells with a low level of hairpin expression and low phenotype penetrance. Taken together, these data suggest that HyFat is required for cell-cell adhesion.

In the body column of *Hydra*, both mitotic spindles [33] and myonemes [11] of epithelial cells are oriented parallel (in the ectoderm), or perpendicular (in the endoderm) to the oral/aboral axis. Epithelial cells of both layers divide symmetrically. As a result each epithelial layer appears as a sheet of a similar-sized cells aligned in the same direction. *In shFat9* and *shFat4* animals epithelial ectodermal cells appeared to be misaligned (Figure 5A). This misalignment was also observed in the endoderm of *shFat9* (Figures 4N,O). Thus HyFat is needed for proper cell alignment.

**Figure 5.**
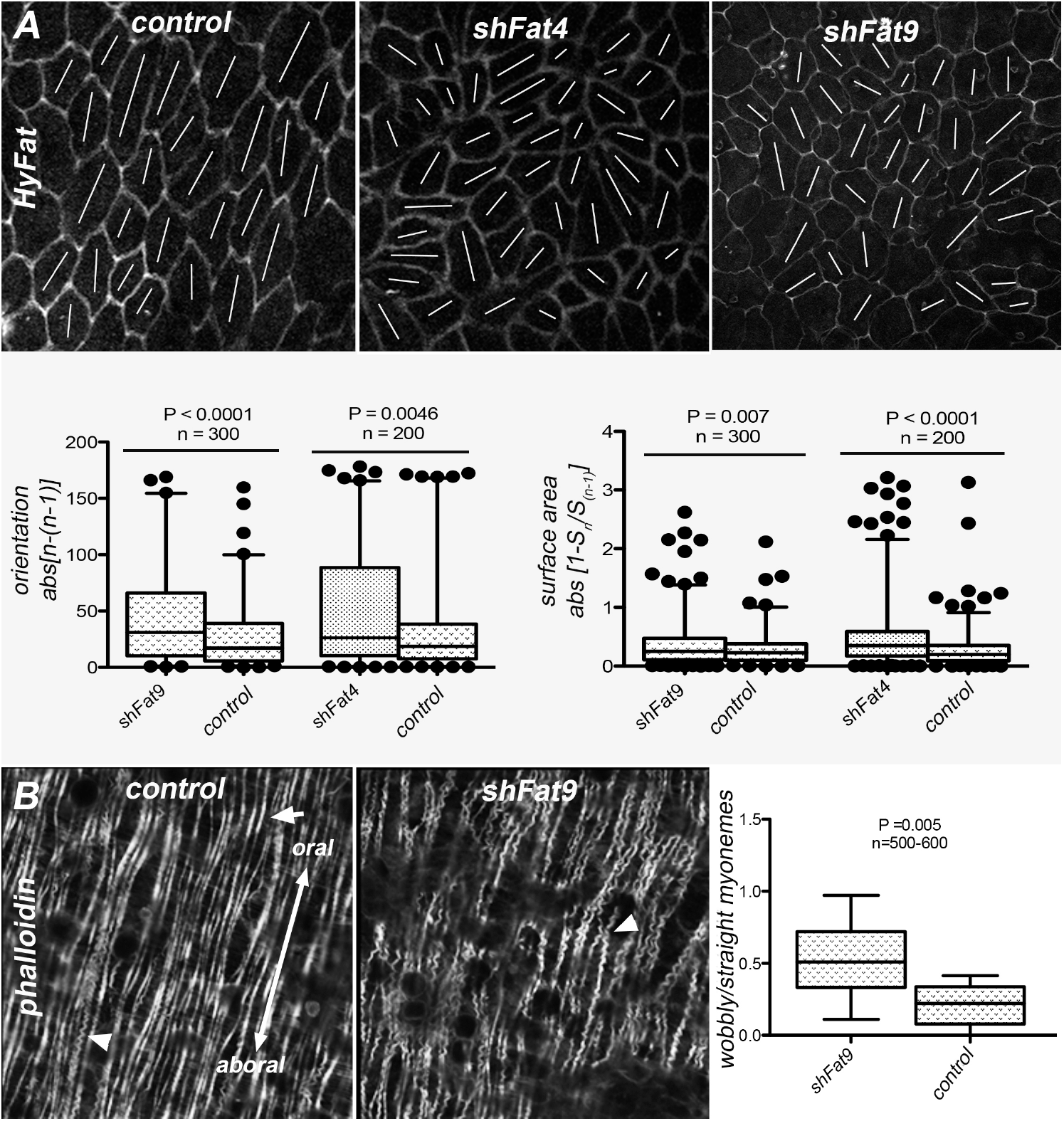
Alignment of ectodermal epithelial cells and morphology of ectodermal actin filaments are affected in *shFat* transgenics. (A) Apical view of ectodermal epithelial cells immunostained with anti-HyFat serum. The graphs show measurements of cell orientation and area using the ‘neighbor’ method. Orientations of individual cells were calculated as an angle between the longest axis of the cell surface and the arbitrary direction using MATLAB. The differences between angles of neighboring cells n-(n-1) were calculated and their absolute values abs[n-(n-1)] were used to measure the degree of misalignment between neighboring cells: the greater the misalignment, the higher the value of abs[n-(n-1)]. To evaluate the differences in the surface area for each pair of neighbor cells, the ratio of the surface area S_n_/ S_n-1_ was calculated, subtracted from 1, and the absolute value abs[1- S_n_/S_n-1_] was used to measure the degree of heterogeneity in the apical surface area. The less similar neighbor cells are to each other, the higher the value of abs[1- S_n_/S_n-1_]. (B) Actin in ectodermal myonemes was visualized with phalloidin; arrow points to the straight and arrowheads to the ‘wobbly’ myonemes, The graph indicates the increased proportion of ‘wobbly’ actin fibers in *shFat9* polyps for n number of myonemes.

In *shFat* knockdowns, epithelial cells also differed from each other in size (Figure 5A). This difference was especially obvious in the basal disk area (Figures 4L,M). We used a ‘neighbor’ method (see Materials and Methods) to measure the degree of cell misalignment and cell size difference. In the absence of cell differentiation, neighboring cells should have a similar size and orientation. In both *shFat9* and *shFat4*, both parameters were significantly different from the control (Figure 5A).

### Myonemes become disorganized and appear “frizzled” upon loss of HyFat

*Hydra* myonemes are muscle processes located on the basal side of epithelial cells and attached to the mesoglea [34]. One of the manifestations of PCP in *Hydra* is a parallel arrangement of the ectodermal muscle processes along the oral/aboral axis. We therefore examined whether downregulation of *HyFat* affects the alignment of ectodermal myonemes. Myonemes in *shFat9 Hydra* were still oriented in the oral-aboral direction. However, in *shFat9*, the actin fibers in the myonemes were less straight than in controls, and had a frizzled appearance (Figure 5B).

### Ectopic expression of the intracellular domain of HyFat affects morphogenesis of ectodermal myonemes

To understand how the intracellular domain of HyFat affects *Hydra* patterning, we generated a transgenic line, *ICD3-GFP*, that constitutively expresses GFP-tagged ICD from HyFat (residues 4100 – 4392) in both ectodermal and endodermal epithelial cells. Mosaic *ICD3-GFP* animals had normal morphology, with ectodermal myonemes oriented parallel to the oral/aboral axis (Figure 6A). The ectodermal myonemes in *ICD3-GFP* animals looked relatively normal, although they were often less homogenous and thicker than in normal polyps (Figure 6A). The number of ectodermal myonemes per unit area was lower in *ICD3-GFP* mutants than in *GFP*-expressing controls and non-transgenic polyps. This could be a result of either fewer processes per cell or shorter processes. Since *GFP*-expressing control animals were fully transgenic and *ICD3-GFP* animals also had many transgenic cells, measuring the length of individual myonemes was difficult. We therefore made chimeric animals by transplanting a transgenic oral half to a non-transgenic aboral half and vice versa (Figure 6B). After 30 - 32 hours, animals were fixed with 4% PFA and immunostained. Ectodermal myonemes of *GFP* and *ICD3-GFP* cells can be seen among the non-transgenic cells in chimeric animals. Figure 7B shows ectodermal myonemes of *GFP* control cells immunostained for GFP and *ICD3-GFP* cells immunostained for HyFat surrounded by the non-transgenic cells. Ectodermal processes of *ICD3-GFP* cells are significantly shorter than those of *GFP* cells (Figure 6C). Thus, ectopic expression of ICD affects the morphology of myonemes.

**Figure 6.**
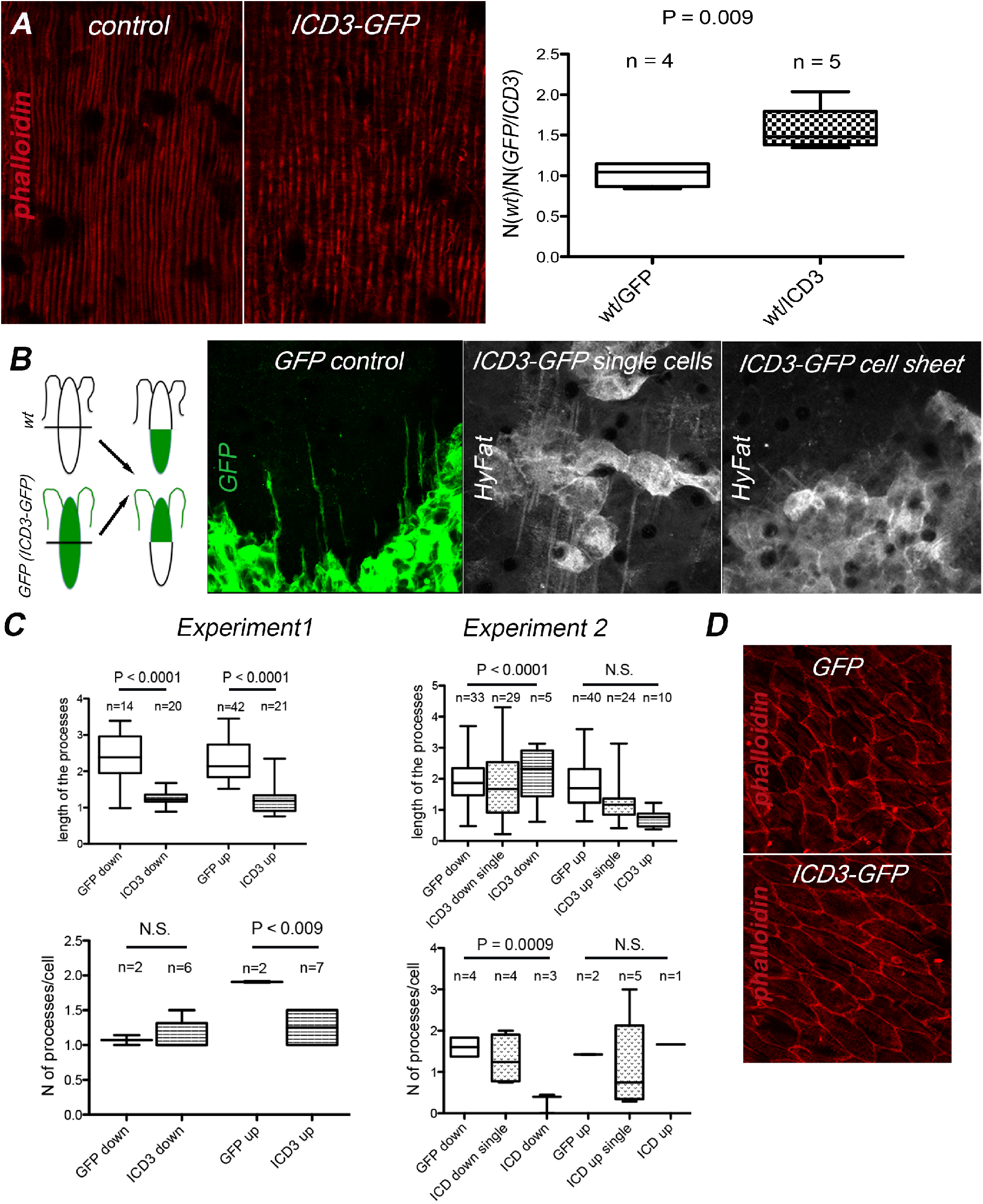
Ectopic expression of the intracellular domain of HyFat affects morphogenesis of ectodermal myonemes. (A) Ectodermal myonemes of *GFP*-expressing and *ICD3* transgenic cells visualized with phalloidin. The graph illustrates the decrease in the number of myonemes in *GFP-ICD3* cells compared to a control polyp; data show the ratio between the number of ectodermal myonemes in the non-transgenic and either *GFP* or *ICD3-GFP* animals determined for n areas; (B) Schematic of *Hydra* transplantations; ectodermal myonemes of *GFP* and *ICD3-GFP* expressing cells in a chimeric *Hydra*; (C) Graphs show the length and number of myonemes/cell determined in Exp. 1 (PFA fixation) and Exp. 2 (Lavdovsky fixation). Myonemes are oriented either towards the hypostome (up) or towards the foot (down); measurements of the lengthes of the myonemes were done using Photoshop and given for the number (n) of myonemes measured; N processes/cell were measured for n number of areas, in each area 2 – 14 transgenic cells were present; (D) Apical view of the ectoderm of *GFP* and *ICD3-GFP* polyps stained with phalloidin.

PFA fixation often causes strong contraction of *Hydra* polyps, therefore we repeated the experiment using a Lavdovsky’s fixative. In this case, significant differences in length were observed only in the myonemes oriented towards the oral end of the animal (Figure 6C). Interestingly, *ICD3-GFP* cells that are single, or within a small group of transgenic cells, have longer myonemes than cells that are a part of an *ICD3-GFP* epithelial sheet. This indicates a non-autonomous effect of ectopic HyFat ICD expression on the morphogenesis of ectodermal myonemes. This is similar to the observation made in *Drosophila* follicle cells, where loss of Fat-like only has effects in large clones of cells [35]. Taken together our results indicate that *HyFat* is involved in regulating the size of the ectodermal epithelial cells as well as the size and local alignment of the ectodermal myonemes

## DISCUSSION

Previous studies have shown that the anthozoan cnidarian *Nematostella* has a complete repertoire of atypical cadherins involved in PCP: Fat, CELSR, Fat-like and Dachsous proteins [17]. However, we found that *Hydra* and other medusozoans lack a Fat homologue and have a single Fat-like cadherin. This finding is surprising since cadherin molecules designated as Fat homologues have been identified in organisms that diverged prior to Cnidaria. These include the sponge *Amphimedon queenslandi* [8], the placozoan *Trichoplax adhaerens* [17], and the choanoflagellate *Monosiga brevicollis* [8]. However, the molecular structure of these Fat proteins, in particular, the number of cadherin repeats, is dramatically different: 14 in *Amphimedon queenslandi*, 37 in *Monosiga brevicollis*, and 9 in *Trichoplax adhaerens*. The composition of the conserved domains in the *Monosiga brevicollis* Fat homologue is also very different from bilaterian Fat cadherins [8]. The similarity of the intracellular domains of the proposed Fat homologues from sponges, placozoans, and choanoflagellates and the cadherins of the Fat family is also very low [8, 17]. It remains to be seen if those organisms have functionally conserved Fat homologues. This leaves the cnidarian class anthozoa as the only non-bilaterian with a clear structural homologue of Fat in its cadherin repertoire. Anthozoans are distinct from other classes of cnidarians in having an internal bilaterality displayed by a pair of directive mesenteries and an asymmetric siphonoglyph, (reviewed in [39]). It is interesting to consider a connection between Fat cadherin and the presence of a second axis of symmetry in anthozoans.

All of the proteins making up the core PCP module have been found in both anthozoan and medusozoan cnidarians [7, 11] as well as in *Trichoplax* and some sponge species [7]. This suggests that the basic PCP machinery was established before the appearance of bilaterians. Addition of a Fat/Ds module to the core complex could have led to a new level of planar cell polarization and appearance of a second axis of symmetry. Notably, the appearance of an M/L axis in anthozoans coincides with the presence of a true Fat cadherin. This intriguing scenario calls for functional studies of *Fat* and *Dachsous* in anthozoans, e.g. the sea anemone *Nematostella*.

A remarkable feature of Fat and Dachsous proteins is their asymmetrical localization in epithelial cells that is crucial for their role in a planar polarization (reviewed in [40]). Planar polarization of Fat-like proteins has also been observed in several cases and is linked to their role in directional cell migration and asymmetric cell morphology [6, 41, 42]. The most explicit example is polarization of *Drosophila* Fat2 at the membranes that are perpendicular to the direction of follicle cell displacement and egg chamber rotation [42, 43].

HyFat is also planar polarized at the epithelial membranes that are perpendicular to the oral/aboral axis of the polyp and the direction of ectodermal epithelial cell displacement. This polarization is observed only in the body column and not in the foot or hypostome. In the foot and the hypostome, as in the body column, HyFat is enriched at the tricellular junctions. Interestingly, the speed of an epithelial cell’s displacement towards the termini is much slower in the foot and the hypostome than in the body column [13]. This dynamic of HyFat localization is reminiscent of Fat2 during egg chamber rotation in *Drosophila*: at early stages when rotation is slow, Fat2 is enriched at the tricellular junctions of the follicle cells, and when rotation enters a fast stage, Fat2 becomes planar polarized [42]. Another interesting similarity between these two homologues is their ability to have a non-autonomous effect, on the morphology of myonemes in the case of HyFat and on the alignment of actin filaments in the case of Fat2 [35].

The complimentary pattern of *HyFat* and *HyDs* expression in *Hydra* testes is intriguing and indicates the possible role of a Fat/Ds heterodimer in the interactions between the maturing spermatogonia and the cells of the ectodermal epithelium that covers them. We were unable to identify a *Hydra* homologue of Four-jointed, a factor that regulates Fat/Ds binding in *Drosophila* and mammals. This suggests that modulation of interactions between HyFat and HyDs might be regulated by a distinct mechanism in *Hydra*.

Fat-like cadherins promote cell migration and polarization through regulation of actin polymerization and microtubule alignment [3–6, 44], via interactions of their intracellular domains with either Ena/VASP in the case of Fat1 and Fat3 or WAVE in the case of Fat2 complexes. The intracellular domain of HyFat contains three proline-rich regions that can potentially interact with the Ena/VASP complex (Figure S1B). Our data show that loss of HyFat interferes with cell shape and cell alignment, possibly through regulation of actin filaments in the ectoderm. Indeed, ectopic expression of HyFat ICD affects the number and the length of ectodermal myonemes. Free HyFat ICD in the cytoplasm could disrupt polarization within the cell and change the morphology of polarized processes. Further functional studies of ICD should reveal the role of HyFat in the regulation of the actin cytoskeleton in *Hydra*.

In summary, our studies show for the first time that the role of atypical cadherins in planar cell polarization and morphogenesis was established before the divergence of cnidarians and bilaterians. The discovery of polarized Fat-like cadherins in early diverging metazoans supports the proposal that subcellular planar polarization of proteins is fundamental to planar polarity of tissues. It remains to be shown whether the true Fat homologue found in anthozoans is linked to bilaterality in this cnidarian class. Testing of this hypothesis could be readily carried out with the tools available in the model anthozoan *Nematostella vectensis*.

## MATERIALS AND METHODS

### Animals, culture conditions, and transplantation experiments

The AEP strain of *Hydra vulgaris* was cultured at 18°C as described previously [45]. Transplantation experiments were done as described in [46].

### In situ hybridization

Whole mount *in situ* hybridization was carried out as described previously [47].The following gene segments were used to make *in situ* probes, numbers indicate positions of nucleotides in the coding sequences: *HhyFat* (12601 – 13178 bp), *HhyDs* (7322 −7621), *myc2* (100 – 605).

### Production of antibodies

Peptides corresponding to the C-terminal portions of HyFat (residues 4191 – 4393) and HyDs (residues 9144 - 9799) were expressed as GST-fusions using the GEX4t-1 vector (Millipore) in *E. coli* strain BL21 and purified on glutathione-agarose (Thermo Scientific). Purified proteins were used to immunize guinea pigs (Cocalico Biologicals) and rats. Polyclonal antibodies were purified by binding to antigen immobilized on glutathione agarose beads and elution with 100 mM glycine (pH 2.5) into 1 M Tris, pH 8.0.

### Immunoblot and Immunofluorescence analysis

For immunoblotting, polyps were dissolved in lysis buffer (5% SDS; 10% glycerol; 60 mM Tris-HCl, pH 6.8) containing 2% beta-mercaptoethanol, boiled for 5 min, chilled on ice for 5 min and electrophoresed in a 3 – 8% Tris-Acetate NuPAGE gel (Novex Life Technologies). Transfer of the proteins onto a nitrocellulose membrane was done in transfer buffer (5% methanol, 25 mM Tris base, 192 mM glycine) overnight at 4° C. The membranes were incubated with antibodies (total anti-HyFat serum was used at 1:1000 dilution, purified anti-HyFat antibodies at 1:100 dilution, anti-actin (clone C4, Millipore) at 1:2000 dilution) in blocking buffer (TBS-tween 0.1% containing 5% powdered milk) O/N at 4°C. After 3×10 min washes with TBS-T, membranes were incubated with HRP-conjugated secondary antibody (GE Healthcare) diluted 1:10000 in blocking buffer for 1 h at RT. Visualization was done by ECL detection (Thermo Scientific). Whole mount immunofluorescence on *Hydra* polyps was done as described previously [48]. Primary antibodies dilutions were as follows: anti-HyFat total serum, 1:1500; affinity purified anti-HyFat, 1:100; anti-HyDs total serum, 1:400. Fluor-conjugated secondary antibodies were used at 1:400 dilution. Alexa555-phalloidin was used at 1:1000 dilution.

### ‘Neighbor’ method to compare apical surface area and orientation of neighboring epithelial cells

Cell outlines were captured using ImageJ. Cell area and orientation were then quantitatively analyzed using a custom image analysis routine in MATLAB. For each pair of neighbor cells, the ratio of the surface area S_n_/ S_n-1_ was calculated, subtracted from 1, and the absolute value abs(1- S_n_/S_n-1_) was used to measure the degree of heterogeneity in the apical surface area. The more similar neighbor cells are to each other, the lower the value of abs(1- S_n_/S_n-1_). A similar approach was used to determine the degree of epithelial cell disorientation. Orientations of individual cells were calculated as an angle between the longest axis of the cell surface and the arbitrary direction using MATLAB. The differences between angles of neighboring cells n-(n-1) were calculated and their absolute values abs[n-(n-1)] were used to measure the degree of misalignment between neighboring cells: the greater the misalignment, the higher the value of abs[n-(n-1)].

### Database search and Phylogenetic Analysis

To identify cnidarian homologues of Fat-like, Dachsous and CELSR proteins we have searched the NCBI (http://www.ncbi.nlm.nih.gov) and NHGRI (https://research.nhgri.nih.gov) databases. As queries, we have used sequences of the intracellular portions of *Nematostella vectensis* Fat-like, Dachsous and CELSR proteins [17]. We have identified *Hydra vulgaris* homologues of Fat-like (XM_012702814.1, former XM_002167158.2), Dachsous (PT_comp25284_c0_seq1) and CELSR (PT_comp27436_c0_seq1), *Alatina alata* partial sequences of Fat-like (GEUJ01029859.1) and Dachsous (GEUJ01029600.1) homologues, *Cassiopea xamachana* partial sequences of Fat-like (OLMO01020275.1) and Dachsous (OLMO01079121.1), *Aurelia aurita* Fat-like homologue (GHAI01167428.1), *Hydractinia symbiolongicarpus* partial Fat-like sequence (GAWH01043253.1). For phylogenetic analysis we have used the following sequences: *Mus musculus* Fat1 (NM_001081286.2), Fat2 (NM_001029988.2), Fat3 (NM_001080814.1), Fat4 (NM_183221.3), CELSR1 (AB028499.1) Dachsous (BC060096.1); *Homo sapience* Fat1 (NM_005245.3) CELSR1 (NM_014246.1), Dachsous1 (AB053446.1) Dachsous2 (NM_001358235.1); *Nematostella vectensis* Fat, Fat-like, Dachsous, CELSR [17]; *Clytia hemisphaerica* Dachsous (JQ438999.1), CELSR (JQ439002.1), Fat-like (JQ439001.1); *Drosophila melanogaster* Fat (NM_058149.4), Kugelei(Fat2) (NM_001300197.1), Dachsous (AAF51468.3), CELSR (AB028498.1), Shotgun (NM_057374.3), Cadherin-N (NM_165227.3); *Branchiostoma floridae* Fat1 (XM_019765978.1), Fat (XM_019791938.1), Dachsous (XM_019791235.1), CELSR [17], cadherin CDH [17]; *Acropora digitifera* Dachsous (XM_015902527.1); *Folsomia candida* Fc1 cadherin (AB190297.1). For generation of the phylogenetic tree, the sequences were aligned using MAFFT (https://www.ebi.ac.uk/Tools/msa/mafft/).

## ACKNOWLEDGEMENTS

The authors are grateful to Bert Hobmayer (University of Innsbruck) for providing the *claudin-GFP* transgenic line and the AEP strain of *Hydra vulgaris*, Jörg Wittlieb (University of Kiel) for generating transgenic lines, and Celina Juliano (University of California, Davis) and Hiroshi Shimizu (KAUST, Thuwal) for providing the AEP *Hydra vulgaris* strain.

## REFERENCES

1. Devenport, D., The cell biology of planar cell polarity. J Cell Biol, 2014. 207(2): p. 171–9.

2. Sharma, P. and H. McNeill, Fat and Dachsous cadherins. Prog Mol Biol Transl Sci, 2013. 116: p. 215–35.

3. Chen, D.Y., et al., Symmetry Breaking in an Edgeless Epithelium by Fat2-Regulated Microtubule Polarity. Cell Rep, 2016. 15(6): p. 1125–33.

4. Squarr, A.J., et al., Fat2 acts through the WAVE regulatory complex to drive collective cell migration during tissue rotation. J Cell Biol, 2016. 212(5): p. 591–603.

5. Moeller, M.J., et al., Protocadherin FAT1 binds Ena/VASP proteins and is necessary for actin dynamics and cell polarization. EMBO J, 2004. 23(19): p. 3769–79.

6. Tanoue, T. and M. Takeichi, Mammalian Fat1 cadherin regulates actin dynamics and cell-cell contact. J Cell Biol, 2004. 165(4): p. 517–28.

7. Schenkelaars, Q., et al., Retracing the path of planar cell polarity. BMC Evol Biol, 2016. 16: p. 69.

8. Abedin, M. and N. King, The premetazoan ancestry of cadherins. Science, 2008. 319(5865): p. 946–8.

9. Momose, T., Y. Kraus, and E. Houliston, A conserved function for Strabismus in establishing planar cell polarity in the ciliated ectoderm during cnidarian larval development. Development, 2012. 139(23): p. 4374–82.

10. Kumburegama, S., et al., Strabismus-mediated primary archenteron invagination is uncoupled from Wnt/beta-catenin-dependent endoderm cell fate specification in Nematostella vectensis (Anthozoa, Cnidaria): Implications for the evolution of gastrulation. Evodevo, 2011. 2(1): p. 2.

11. Philipp, I., et al., Wnt/beta-catenin and noncanonical Wnt signaling interact in tissue evagination in the simple eumetazoan Hydra. Proc Natl Acad Sci U S A, 2009. 106(11): p. 4290–5.

12. Bode, H.R., The interstitial cell lineage of hydra: a stem cell system that arose early in evolution. J Cell Sci, 1996. 109 (Pt 6): p. 1155–64.

13. Campbell, R.D., Tissue dynamics of steady state growth in Hydra littoralis. II. Patterns of tissue movement. J Morphol, 1967. 121(1): p. 19–28.

14. Bode, H.R., Axial patterning in hydra. Cold Spring Harb Perspect Biol, 2009. 1(1): p. a000463.

15. Niebuhr, K., et al., A novel proline-rich motif present in ActA of Listeria monocytogenes and cytoskeletal proteins is the ligand for the EVH1 domain, a protein module present in the Ena/VASP family. EMBO J, 1997. 16(17): p. 5433–44.

16. Hulpiau, P. and F. van Roy, Molecular evolution of the cadherin superfamily. Int J Biochem Cell Biol, 2009. 41(2): p. 349–69.

17. Hulpiau, P. and F. van Roy, New insights into the evolution of metazoan cadherins. Mol Biol Evol, 2011. 28(1): p. 647–57.

18. Usui, T., et al., Flamingo, a seven-pass transmembrane cadherin, regulates planar cell polarity under the control of Frizzled. Cell, 1999. 98(5): p. 585–95.

19. Morris, L.G., et al., Recurrent somatic mutation of FAT1 in multiple human cancers leads to aberrant Wnt activation. Nat Genet, 2013. 45(3): p. 253–61.

20. Cho, E. and K.D. Irvine, Action of fat, four-jointed, dachsous and dachs in distal-to-proximal wing signaling. Development, 2004. 131(18): p. 4489–500.

21. Otto, J.J. and R.D. Campbell, Budding in Hydra attenuata: bud stages and fate map. J Exp Zool, 1977. 200(3): p. 417–28.

22. Kuznetsov, S., M. Lyanguzowa, and T.C. Bosch, Role of epithelial cells and programmed cell death in Hydra spermatogenesis. Zoology (Jena), 2001. 104(1): p. 25–31.

23. Littlefield, C.L. and H.R. Bode, Germ cells in Hydra oligactis males. II. Evidence for a subpopulation of interstitial stem cells whose differentiation is limited to sperm production. Dev Biol, 1986. 116(2): p. 381–6.

24. Zakaria, S., et al., Regulation of neuronal migration by Dchs1-Fat4 planar cell polarity. Curr Biol, 2014. 24(14): p. 1620–1627.

25. O’Donovan, P. and M. Abraham, Somatic tissue-male germ cell barrier in Hydra viridis (hydrozoa, coelenterata). J Morphol, 1988. 198(2): p. 179–188.

26. Sadeqzadeh, E., et al., Dual processing of FAT1 cadherin protein by human melanoma cells generates distinct protein products. J Biol Chem, 2011. 286(32): p. 28181–91.

27. Sing, A., et al., The atypical cadherin fat directly regulates mitochondrial function and metabolic state. Cell, 2014. 158(6): p. 1293–1308.

28. Aufschnaiter, R., et al., Apical and basal epitheliomuscular F-actin dynamics during Hydra bud evagination. Biol Open, 2017. 6(8): p. 1137–1148.

29. Boehm, A.M., et al., FoxO is a critical regulator of stem cell maintenance in immortal Hydra. Proc Natl Acad Sci U S A, 2012. 109(48): p. 19697–702.

30. Klimovich, A., et al., Non-senescent Hydra tolerates severe disturbances in the nuclear lamina. Aging (Albany NY), 2018. 10(5): p. 951–972.

31. Wittlieb, J., et al., Transgenic Hydra allow in vivo tracking of individual stem cells during morphogenesis. Proc Natl Acad Sci U S A, 2006. 103(16): p. 6208–11.

32. Muller, W.A., Ectopic head and foot formation in Hydra: diacylglycerol-induced increase in positional value and assistance of the head in foot formation. Differentiation, 1990. 42(3): p. 131–43.

33. Shimizu, H., P.M. Bode, and H.R. Bode, Patterns of oriented cell division during the steady-state morphogenesis of the body column in hydra. Dev Dyn, 1995. 204(4): p. 349–57.

34. West, D.L., The epitheliomuscular cell of hydra: its fine structure, three-dimensional architecture and relation to morphogenesis. Tissue Cell, 1978. 10(4): p. 629–46.

35. Viktorinova, I., et al., Modelling planar polarity of epithelia: the role of signal relay in collective cell polarization. J R Soc Interface, 2011. 8(60): p. 1059–63.

36. Browne, E.N., The production of new hydrants in hydra by the insertion of small grafts. Journal of Experimental Zoology, 1909. 7: p. 1 – 37.

37. Hobmayer, B., et al., WNT signalling molecules act in axis formation in the diploblastic metazoan Hydra. Nature, 2000. 407(6801): p. 186–9.

38. Broun, M., et al., Formation of the head organizer in hydra involves the canonical Wnt pathway. Development, 2005. 132(12): p. 2907–16.

39. Berking, S., Generation of bilateral symmetry in Anthozoa: a model. J Theor Biol, 2007. 246(3): p. 477–90.

40. Blair, S. and H. McNeill, Big roles for Fat cadherins. Curr Opin Cell Biol, 2018. 51: p. 73–80.

41. Krol, A., S.J. Henle, and L.V. Goodrich, Fat3 and Ena/VASP proteins influence the emergence of asymmetric cell morphology in the developing retina. Development, 2016. 143(12): p. 2172–82.

42. Viktorinova, I., et al., The cadherin Fat2 is required for planar cell polarity in the Drosophila ovary. Development, 2009. 136(24): p. 4123–32.

43. Barlan, K., M. Cetera, and S. Horne-Badovinac, Fat2 and Lar Define a Basally Localized Planar Signaling System Controlling Collective Cell Migration. Dev Cell, 2017. 40(5): p. 467–477 e5.

44. Cheng, H., et al., Disparate Regulatory Mechanisms Control Fat3 and P75NTR Protein Transport through a Conserved Kif5-Interaction Domain. PLoS One, 2016. 11(10): p. e0165519.

45. Smith, K.M., et al., CnOtx, a member of the Otx gene family, has a role in cell movement in hydra. Dev Biol, 1999. 212(2): p. 392–404.

46. Broun, M. and H.R. Bode, Characterization of the head organizer in hydra. Development, 2002. 129(4): p. 875–84.

47. Grens, A., et al., CnNK-2, an NK-2 homeobox gene, has a role in patterning the basal end of the axis in hydra. Dev Biol, 1996. 180(2): p. 473–88.

48. Brooun, M., et al., Organizer formation in Hydra is disrupted by thalidomide treatment. Dev Biol, 2013. 378(1): p. 51–63.

