## Supplemental Material for "Ancestral role of Fat-like cadherins in planar cell polarity"

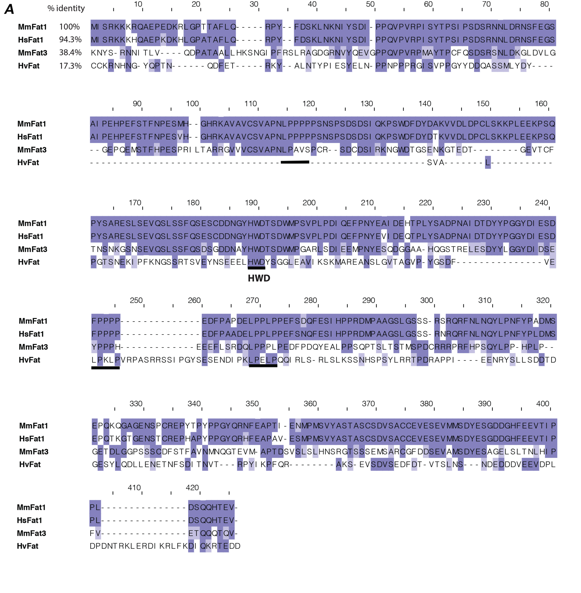

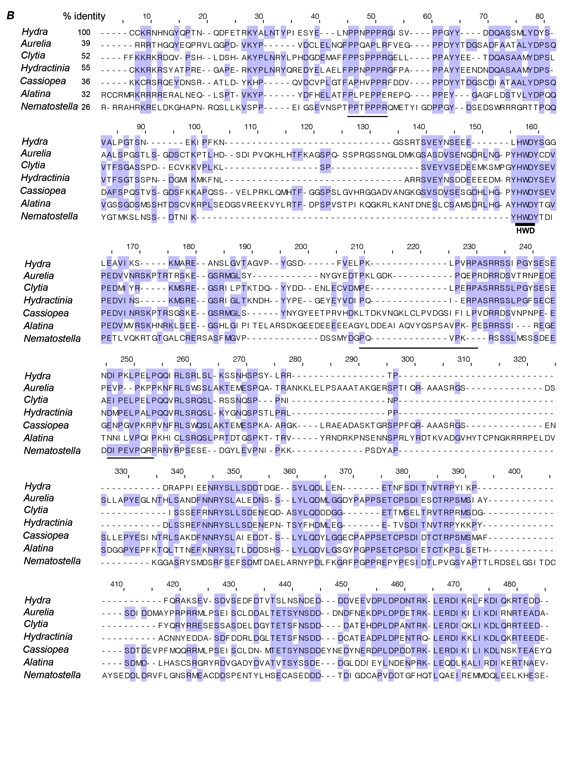

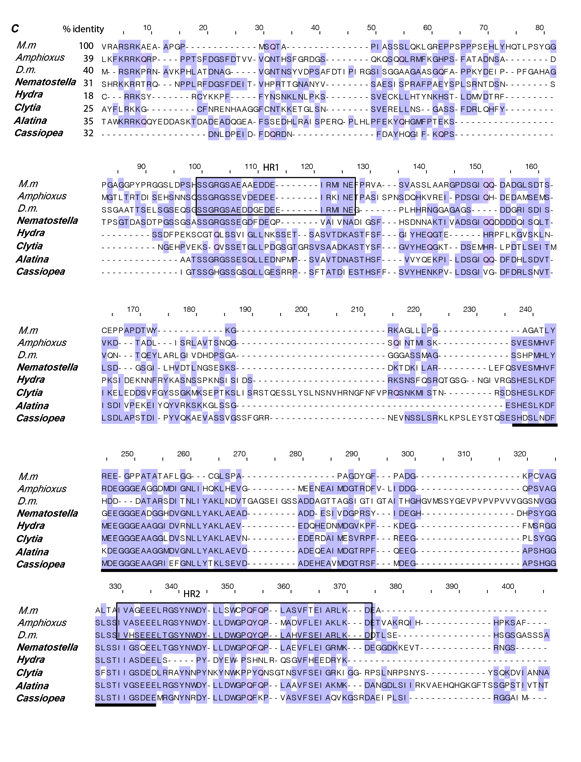

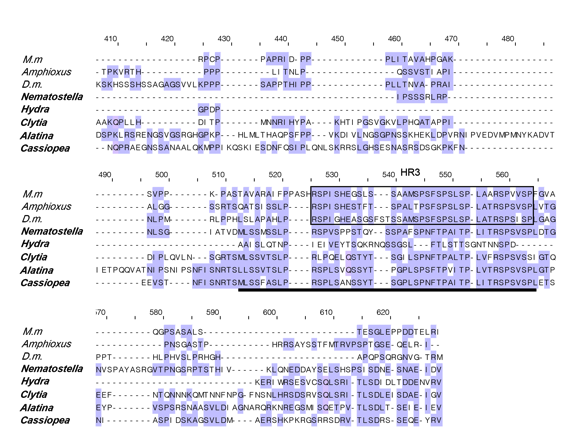

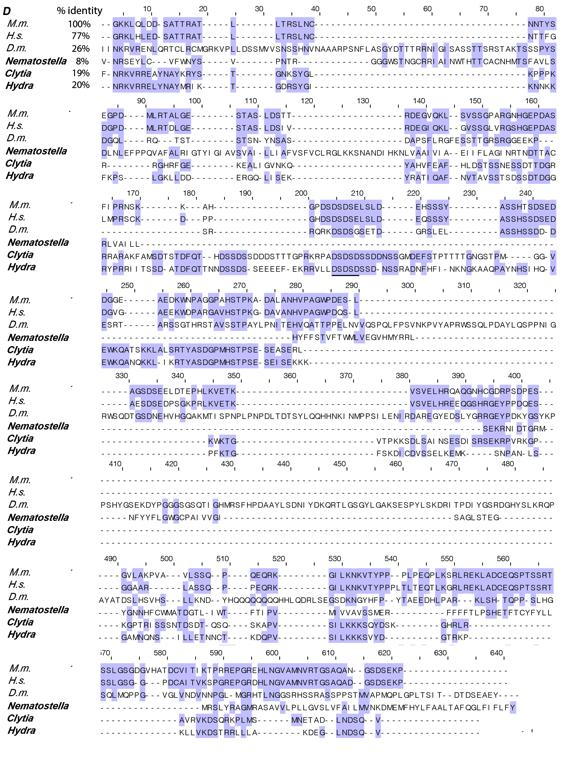

**Figure S1** **Amino acid sequence alignments of ICDs of atypical cadherins.** Alignments were done using MAFFT (https://www.ebi.ac.uk/Tools/msa/mafft/) and visualized with PFAAT (http://pfaat.sourceforge.net). Identical residues are highlighted in blue. (A) Mammalian Fat-like and predicted *Hydra* Fat-like proteins show low similarity. *Hydra* proline-rich, presumable EVH1-like domains are underlined; (B) Cnidarian Fat-like sequences show significant similarity with the exception of *Nematostella*. Putative proline-rich EVH1-like domains are underlined; (C) Weak similarity between cnidarian Ds sequences and other Ds homologues; homologous regions within cnidarians are underlined; homologous regions of chordate homologues (HR) are in boxes. (D) CELSR homologues.

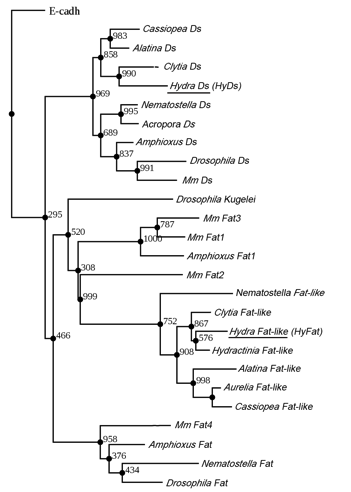

**Figure S2** **Phylogenetic tree of Fat, Fat-like, and Dachsous subfamilies based on ICD sequences.** Maximum likelihood analysis (1000 bootstrap replicates, bootstrap values are indicated for each node) supports the clustering of the predicted *Hydra* homologues (underlined) with the Fat-like and Dachsous cadherin subfamilies.

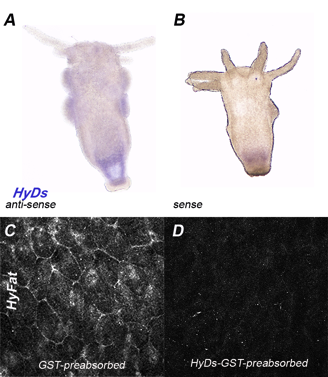

**Figure S3** **HyDs is expressed in *Hydra* epithelial cells.** (A,B) Expression of *HyDs* in *Hydra* epithelial cells determined by in situ hybridization; (C,D) Apical view of ectodermal epithelial cells of *Hydra* immunostained with HyDs anti-serum (1:200) preabsorbed with GST (C) or HyDs-GST antigen (D).

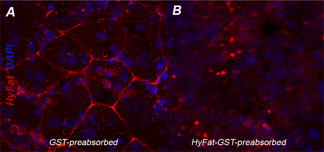

**Figure S4** **Preabsorbtion with antigen confirms the specificity of the anti-HyFat antiserum.** Apical view of ectodermal epithelial cells of *Hydra* immunostained with HyFat anti-serum (1:1000) that was preabsorbed with (A) GST protein or (B) HyFat-GST antigen.

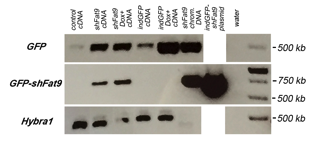

**Figure S5** **Evaluation of *shFat9* hairpin expression in transgenic *shFat9* polyps by RT-PCR.** PCR with primers for the *HyBra1* gene was performed as a control for cDNA preparation: the 500 bp fragment cannot be amplified from genomic DNA because sequences of the forward and reverse primers are separated by an intron. PCR was performed for 35 cycles and the primers were as follows: for *GFP-shFat9* the forward primer was 4000 – 4024 bp of pIndGFP (3’ end of *GFP* gene), the reverse primer to was 13143 – 13178 bp of *HyFat* transcript; for the GFP fragment the forward primer was 3400 – 3426 bp and the reverse primer was 3899 – 3920 bp of pIndGFP; for *HyBra1* the forward primer was 798 – 827 bp and the reverse primer was 1308 – 1334 bp of *HyBra1* transcript (AY366371.1).

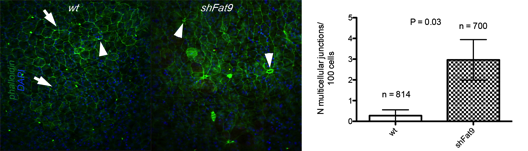

**Figure S6** **Multicellular junctions in the *shFat9* ectoderm.** Apical view of the ectoderm of non-transgenic and *shFat9* polyps stained with phalloidin. Arrows indicate three- and four-cell junctions, arrowheads indicate more than four cell junctions. The graph shows the number of multicellular junctions per 100 ectodermal epithelial cells, the data represent mean with SEM.
